# CRISPR-mediated isogenic cell-SELEX approach for generating highly specific aptamers against native membrane proteins

**DOI:** 10.1101/2020.02.17.949768

**Authors:** Jonah C. Rosch, Emma H. Neal, Daniel A. Balikov, Mohsin Rahim, Ethan S. Lippmann

**Author notes:** Corresponding author: Ethan S. Lippmann PMB 351604, 2301 Vanderbilt Place, Nashville, TN 37235-1604.

## Abstract

**Introduction:** The generation of affinity reagents that bind native membrane proteins with high specificity remains challenging. Most *in vitro* selection paradigms utilize different cell types for positive and negative rounds of selection (where the positive selection is against a cell that expresses the desired membrane protein and the negative selection is against a cell that lacks the protein). However, this strategy can yield affinity reagents that bind unintended membrane proteins on the target cells. To address this issue, we developed a systematic evolution of ligands by exponential enrichment (SELEX) scheme that utilizes isogenic pairs of cells generated via CRISPR techniques.

**Methods:** Using a Caco-2 epithelial cell line with constitutive Cas9 expression, we knocked out the *SLC2A1* gene (encoding the GLUT1 glucose transporter) via lipofection with synthetic gRNAs. Cell-SELEX rounds were carried out against wild-type and GLUT1-null cells using a single-strand DNA (ssDNA) library. Next-generation sequencing (NGS) was used to quantify enrichment of prospective binders to the wild-type cells.

**Results:** 10 rounds of cell-SELEX were conducted via simultaneous exposure of ssDNA pools to wild-type and GLUT1-null Caco-2 cells under continuous perfusion. The top binders identified from NGS were validated by flow cytometry and immunostaining for their specificity to the GLUT1 receptor.

**Conclusions:** Our data indicate that highly specific aptamers can be isolated with a SELEX strategy that utilizes isogenic cell lines. This approach should be broadly useful for generating affinity reagents that selectively bind to membrane proteins in their native conformations on the cell surface.

## Introduction

To support basic research applications and clinical translation, there is an ever-increasing demand for the development of high-specificity affinity reagents for biological targets [1],[2]. While *in vitro* selection strategies have been used to develop many types of affinity reagents (including nucleic acid aptamers, peptides, and proteins) for a wide range of biological targets, the ability to generate highly specific binders to native membrane proteins remains challenging. Selection strategies for membrane proteins are often facilitated by the use of recombinant extracellular domains, but this approach has drawbacks because the recombinant domain may not properly fold due to the lack of a transmembrane anchor or include appropriate post-translational modifications; both of these issues can affect the ability of the binding reagent to selectively recognize the membrane protein when it is properly expressed in a cell lipid bilayer. Attempts to purify cell-surface targets into artificial support systems like nanodiscs can similarly lead to a loss of endogenous conformations and appropriate protein-protein interactions, while also imposing yield limitations. For these reasons, many selection strategies utilize whole cells to preserve membrane protein integrity. For example, nucleic acid libraries can be used to select aptamers against whole cells via standard SELEX (Systematic Evolution of Ligands through Exponential Enrichment) approaches [3], whereas display techniques can be used to select peptide/protein-based affinity reagents against whole cells [4], [5]. These strategies have yielded binding reagents against various membrane proteins, including but not limited to, the glucagon receptor [6], EGFR [7], TGFBRIII [8], and PDGFRβ [9].

However, whole cell selections still have drawbacks. For example, most *in vitro* selections utilize different cell types for positive and negative rounds of selection. In this manner, the positive selection is performed on a cell type that expresses the desired membrane target and the negative selection is performed against a cell type that lacks the protein [10], [11]. This strategy can lead to the isolation of affinity reagents that bind unintended membrane proteins on the surface of cells. Attempts to use isogenic cells in the positive and negative selection steps have involved overexpressing the target of interest in a cell type and counterselecting with the parental cell line [12], [13], but this method can still lack sufficient counterselection stringency to yield aptamers with high specificity. Others have attempted sequential selections on recombinant, truncated extracellular motifs followed by whole cell biopanning [14], [15]. This approach has been shown to help improve specificity during the selection process but also reintroduces the cumbersome use of purified protein. Thus, there is substantial room for improvement in whole cell selection workflows for generating affinity reagents.

In this current study, we focus on the development of a novel SELEX approach that uses isogenic cell pairs to generate aptamers against membrane proteins. Aptamers are oligonucleotide-based affinity reagents that have been selected to bind to a variety of targets, including small molecules, proteins, and cell surface receptors. Properly selected aptamers are able to bind to their target with high affinity and specificity, with the added benefits of inexpensive and reproducible chemical synthesis, facile chemical modification, and low immunogenicity [16]. The selection of aptamers by SELEX is usually performed with isolated targets, with 71% of published targets for purified recombinant protein and 19% for synthesized small molecules [17]. In contrast, less than 10% of published aptamer papers have sought to identify aptamers for specific targets on the cell surface, likely due to the difficulties mentioned above. To overcome the previously mentioned issues with whole cell-SELEX, we developed a strategy that utilizes isogenic pairs of cells generated with clustered regularly interspaced short palindromic repeats (CRISPR) techniques (**Fig 1**). In this approach, cell types are used in the positive and negative selection rounds that are prospectively identical except for the one cell surface protein of interest that has been knocked out, which we hypothesized would drive the specificity of the selection. The generation of knockouts with modern CRISPR technologies helps ensure a high specificity of indel formation in the desired gene of interest. For this proof-of-concept endeavor, we chose to generate aptamers against glucose transporter 1 (GLUT1) by knocking out the *SLC2A1* gene to engineer a null cell as an isogenic pair. GLUT1 is implicated in several diseases including Glucose Transporter Type 1 Deficiency Syndrome [18], Alzheimer’s Disease [19], and brain microvasculature defects [20]. Several well-characterized antibodies exist for GLUT1, which is useful for direct comparisons to selected aptamers; however, very few of these antibodies target the extracellular domain of GLUT1, indicating that a high-fidelity aptamer would have utility for research applications. Additionally, GLUT1 is a member of 14 GLUT family members that are structurally similar, and GLUT1 is particularly homologous to GLUT2-4 [21]; likewise, GLUT1 has minimal extracellular exposure, with five short extracellular loops (<15 amino acids each) and one longer loop of 33 amino acids [22]. Thus, GLUT1 is an ideal test case for this study because: (1) it shares substantial structural homology with other transporters in the same family, permitting assessments of aptamer specificity after selection, and (2) its structure is relatively difficult to target extracellularly with an affinity reagent compared to other ligands such as single-pass transmembrane proteins that have a longer extracellular domain. This new selection strategy, deemed CRISPR-mediated isogenic cell-SELEX, successfully generated an aptamer that specifically binds to the GLUT1 transporter on multiple cell types and in human brain tissue. We expect that this strategy will help researchers overcome many difficulties involved with cell-SELEX and identify improved affinity reagents with high specificity for cell membrane proteins.

**Figure 1.**
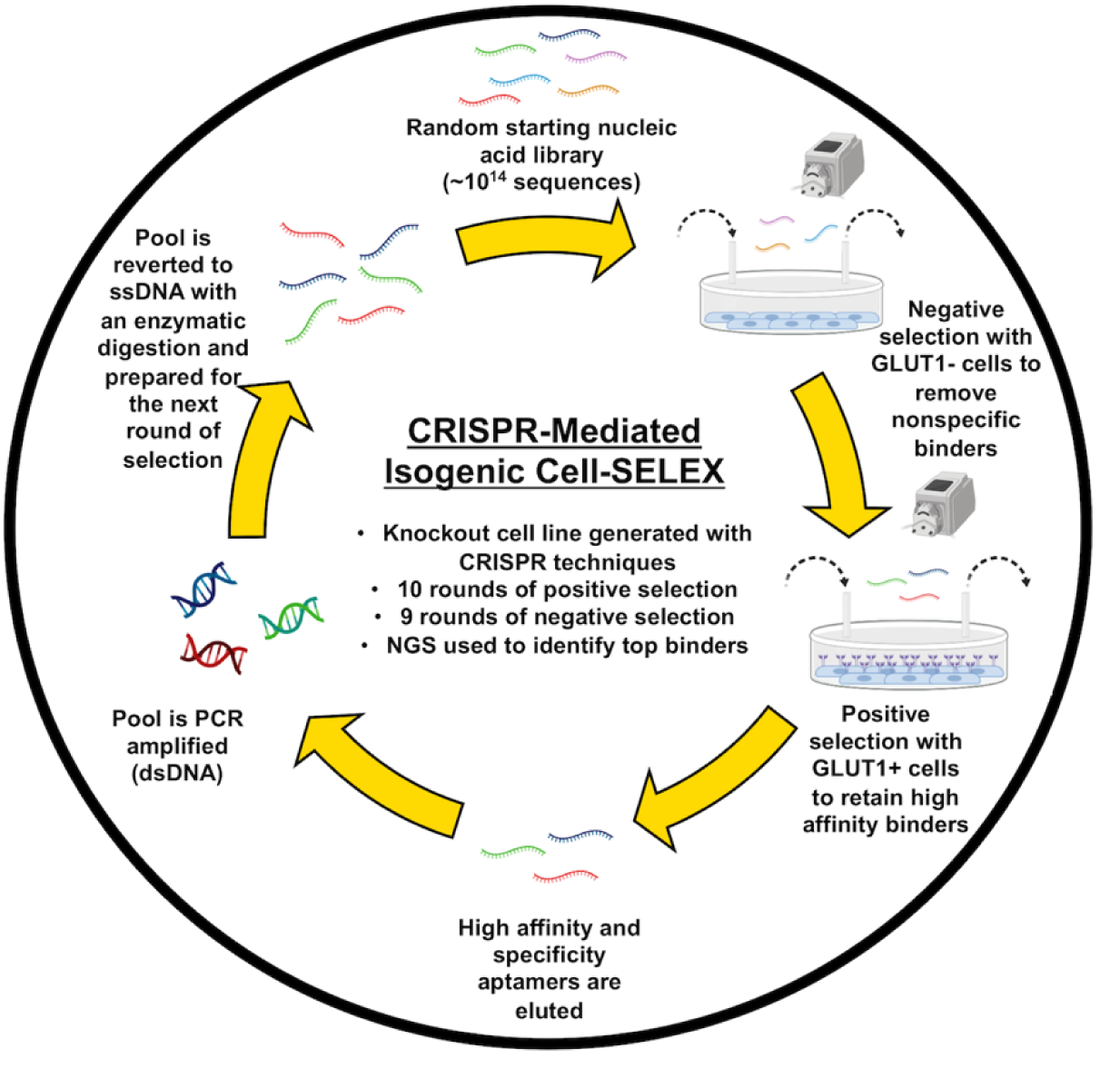
Overview of CRISPR-mediated isogenic cell-SELEX. A random ssDNA library is perfused over null cells, which do not express the protein of interest, as the negative selection. The library is simultaneously perfused over wild-type cells, which express the protein of interest, as the positive selection. Aptamers are eluted from the wild-type cells, PCR amplified, and regenerated to ssDNA by enzymatic digestion to be introduced into the next round of selection. For this study, ten rounds of selection were performed and analyzed with next generation sequencing.

## Materials and Methods

### Materials

All benchtop laboratory materials were purchased from Fisher Scientific and all chemicals were purchased from Sigma Aldrich, unless otherwise stated.

### Cell lines and cell culture

Human Caco-2, MDA-MB-231, and HEK-293 cells were cultured in high glucose DMEM (Corning; 10-0130CV), with sodium pyruvate and L-glutamine, 10% heat inactivated fetal bovine serum (FBS) (Gibco, 26140079), 1X MEM non-essential amino acid solution (Sigma, M7145), and 1% penicillin-streptomycin (Gibco, P0781). FBS was heat-inactivated by heating at 56°C for 30 min followed by cooling on ice. Caco-2 cells were cultured on 0.1% gelatin (Sigma, G1890). Human brain microvascular endothelial cells were differentiated from induced pluripotent stem cells and purified according to previous protocols routinely used in our lab [23], [24]. All cells were cultured at 37°C in a 5% CO_2_ humid atmosphere.

### Strategy for developing GLUT1-null Caco-2 cells

To incorporate Cas9 expression into the Caco-2 cell line, cells were washed twice with DPBS, dissociated with TrypLE Select (Gibco, 1253029), and collected as a single-cell suspension. Based on cell density, this suspension was transduced with a multiplicity of infection (MOI) of 0.3 with Edit-R lentiviral mKate2-tagged, constitutively expressed Cas9 nuclease under the hEF1α promoter (Dharmacon, CAS11229) in transduction medium containing 5 μg/mL polybrene (EMD Millipore, TR-1003-G). Complete growth medium was added to the transduced cells at a 3:1 dilution after 5 hours, and medium was changed every 48 hours prior to fluorescence activated cell sorting (FACS). Once ready for sorting, cells were dissociated and resuspended in phenol-red free DMEM medium (Gibco, 21063-029) supplemented with 10 μM Y27632 dihydrochloride (Tocris, 1254) and antibiotic-antimycotic (Gibco, 15240062). Cells were clonally sorted onto 96-well plates using a 5-laser FACS Aria III (BD Biosciences) with a 100 μm nozzle. Sorted cells were collected in sterile-filtered conditioned medium supplemented with 10μM Y27632, 10 mM HEPES (Gibco, 15630080), and 1X penicillin-streptomycin. Clones were supplemented with an additional 100 μL of complete growth medium per well approximately 24 hours after sorting. Medium was changed with complete growth media every 48 hours after. Cas9 expression of the clones was validated with western blot and immunofluorescent imaging with a Cas9 antibody (Abcam, ab20444) and an occludin antibody (Thermo, 33-1500), similar to methods described below. Clone C6 was selected for further analysis and experimental use.

To generate knockouts, *SLC2A1* crRNA (CM-007509-01, CM-007509-02, CM-007509-03, CM-007509-04) and synthetic tracrRNA (U-002005-05) were ordered from Horizon Discovery and resuspended according to the company’s instructions. Knockout rounds were performed by incubating precomplexed 25 nM crRNA, 25 nM tracrRNA, and Dharmafect 1 reagent (T-2001-01) with Caco-2 cells in antibiotic-free medium for 48 hours at 37°C. After 48 hours, the cells were switched to normal medium, allowed to become confluent, and then subjected to another round of gRNA delivery. After three transfections, the Caco-2 cells were re-seeded at a low density on a 150 mm dish (Fisher, FB0875714) and clonally expanded. 20 cell colonies were picked, expanded, and prospectively analyzed for GLUT1 expression by western blot (data not shown). A single clone was then used for full validation of GLUT1 knockout.

### Validating GLUT1 knockout in Caco-2 cells

GLUT1 knockout in the Caco-2 clonal cell line was determined with three analysis methods (western blot, immunofluorescent imaging, and Sanger sequencing followed by indel analysis). For western blotting, total protein was collected from cell lysate in RIPA buffer (Sigma, R0278) containing 1% phosphatase inhibitor cocktail 3 (Sigma, P0044) and 1% protease inhibitor cocktail (Sigma, P8340), followed by incubation on ice for 30 minutes with occasional pipetting. The RIPA mixture was centrifuged at 12,000xg at 4°C for 15 minutes to isolate protein. Protein concentration was measured with a standard BCA assay (Thermo Fisher, 23225). 10 μg of protein for each condition was run on a 4-20% polyacrylamide gel (Bio-Rad, 5671094), transferred to nitrocellulose membranes (Thermo Fisher, IB23001) using the iBlot2 Gel Transfer Device (Thermo Fisher), blocked for 1 hour with Intercept TBS Blocking Buffer (Li-Cor, 927-600001), and incubated with primary antibodies overnight at 4°C (GLUT1: Abcam, 15309, 1:1000 dilution; GAPDH: CST, D16H11, 1:1000 dilution). Membranes were washed with TBS-T (TBS Buffer with 0.05% Tween-20), incubated with IRDye 800CW goat anti-rabbit secondary antibody (Li-Cor, 926-32211) for 1 hour, washed 3 times with TBS-T, and imaged on an Odyssey Fc Imager (Li-Cor).

For immunofluorescent imaging of cells, cells were washed twice with PBS and fixed in 4% paraformaldehyde (Thermo Fisher, J19942-K2) for 10 minutes. Fixed cells were washed three times, blocked with PBS containing 5% donkey serum and 0.3% Triton-X-100 for 1 hour, and incubated with primary GLUT1 antibody (Abcam, 15309, 0.5 μg/mL) overnight at 4°C. Cells were washed 5 times with PBS, incubated with Alexa Fluor 488 donkey anti-rabbit secondary antibody (Thermo Fisher, A-21206) in the dark for 1 hour, incubated with DAPI nuclear stain (Thermo Fisher, D1306, 1:10000 dilution) for 10 minutes, washed 3 times, and imaged on a Leica DMi8 fluorescence microscope. Cell images were analyzed on FIJI.

For Tracking of Indels by Decomposition (TIDE) [25] analysis, genomic DNA primers flanking gRNA cute sites in the *SLC2A1* gene were designed using the NCBI Primer-BLAST website. The top primers for each cut site were ordered from Integrated DNA Technologies (IDT). Genomic DNA was extracted from wild-type and GLUT1-null Caco-2 cells with QuickExtract DNA Extraction Solution (Lucigen, SS000035-D2) and regions of interest were PCR amplified with the designed primers and Taq polymerase (NEB, M0273S). Amplified DNA was cleaned up with the Monarch DNA Gel Extraction Kit (NEB, T1020S) and samples were submitted to Genewiz for standard Sanger sequencing analysis. Sanger sequencing files for each cut site were analyzed on the TIDE online web tool to determine the indel formation percentage between the wild-type and GLUT1-null cell lines.

### DNA aptamer library, primers, and buffers used for SELEX

The ssDNA aptamer library was synthesized and HPLC-purified by IDT, with 40 nucleotide random bases flanked by 20 nucleotide primer ends required to perform PCR amplification (forward fixed region: TCGCACATTCCGCTTCTACC, reverse fixed region: CGTAAGTCCGTGTGTGCGAA). The starting library was designed with a A:C:G:T molar ratio of 3:3:2:2.4 to adjust for equimolar amounts of nucleotide incorporation. Primers used included: forward primer, TCGCACATTCCGCTTCTACC; 5’-biotinylated forward primer, /5bio/TCGCACATTCCGCTTCTACC; 5’-FAM-labeled forward primer, /5FAM/TCGCACATTCCGCTTCTACC; reverse primer, TTCGCACACACGGACTTACG; 5’-phosphorylated reverse primer, /5Phos/TTCGCACACACGGACTTACG. Wash buffer was prepared with 5 mM MgCl_2_ (Thermo Fisher, AM9530G) and 4.5 g/L glucose (RPI, G32040) in DPBS. Binding buffer was prepared with 100 mg/mL of yeast tRNA (Thermo Fisher, AM7119) and 1 mg/mL of bovine serum albumin (RPI, A30075).

### Cell-SELEX procedure

A recently published protocol for performing SELEX on isolated protein under dynamic flow conditions was used as a starting point for developing this CRISPR-mediated isogenic cell-SELEX protocol [26]. Four nanomoles of ssDNA aptamer library was diluted in binding buffer, heated at 95°C for 10 minutes, cooled on ice for 10 minutes, and then loaded into tubing of a peristaltic pump (Fisher Scientific, 13-310-661). For the first round of cell-SELEX, wild-type Caco-2 cells were seeded into a 0.1% gelatin coated 150 mm cell culture dish (Fisher Scientific, 12565100) and grown until just fully confluent. The initial ssDNA library was circulated over the wild-type Caco-2 cells for 1 hour at 4°C. After positive incubation, washing was conducted by pumping washing buffer over the cells for 15 minutes to remove weakly bound aptamers. After washing, bound aptamers were eluted by scraping the cells into ultrapure water and heating for 10 minutes at 95°C. The mixture was spun at 13,000xg for 5 minutes and the supernatant containing the unbound aptamers was collected.

The Round 1 eluted pool was PCR amplified for 10 cycles with GoTaq Hot Start Polymerase (Promega, M5001) (1X Colorless GoTaq Flexi Buffer, 0.2mM each dNTP, 0.2 uM forward and reverse primers, and 1.25 U Polymerase) [cycling conditions: 95°C for 2 minutes, 10 cycles of 95°C for 30 seconds, 56°C for 30 seconds, 72°C for 1 minutes]. A pilot PCR amplification was performed to determine the optimal PCR cycle conditions for scaleup by testing amplification cycles of 8, 10, 12, 14, and 16. The bands from this pilot test were checked on a 3% agarose gel to ensure no smearing or higher molecular weight byproduct formation (data not shown). The optimized PCR was then performed with unlabeled forward primers and phosphate-labeled reverse primers. Double-stranded DNA (dsDNA) was reverted to ssDNA by incubating with 5U of λ-exonuclease (NEB, M0262S) at 37°C for 1 hour, followed by heat inactivation at 75°C for 10 minutes in a standard PCR thermocycler. The single-strand product was purified through phenol/chloroform/isomayl alcohol extraction (Thermo Fisher, P3803) and ethanol precipitated. ssDNA was resuspended in binding buffer before being used in the next round of selection.

After Round 1, a negative selection step was incorporated into the procedure. In Rounds 2-10, GLUT1-null Caco-2 cells were seeded into a 0.1% gelatin coated 35 mm cell culture treated dish (Fisher, 1256590) overnight and GLUT1-positive wild-type Caco-2 cells were seeded into a 0.1% gelatin coated 100 mm cell culture treated dish (Fisher, FB012924) overnight. The amplified Round 1 ssDNA pool was continuously recirculated over each cell type in series for 1 hour at 4°C, followed by washing with wash buffer for 15 minutes. Following the primary and negative selection step, bound ssDNA was again eluted, PCR-amplified to dsDNA and subsequently restored to ssDNA in a similar manner as described above. To increase selection pressure, the plate size of the negative cells was increased from 35 mm to 150 mm, the plate size of the wild-type cells was decreased from 150 mm to 100 mm (thus manipulating the ratio of positive to negative cells), and the time of washing increased from 15 minutes to 30 minutes by Round 6. For next-generation sequencing (NGS), dsDNA from wild-type cell samples was saved from Rounds 2, 4, 6, 8, and 10 and dsDNA from knockout-cell samples was saved from Rounds 6 and 10.

### Pool affinity characterization

After six rounds of selection, a flow cytometry assay was used to qualitatively determine the affinity of the pool toward the wild-type and GLUT1-null Caco-2 cells. A Guava EasyCyte (Luminex) was used for all flow cytometry experiments. The Round 6 pool and starting library were PCR-amplified with FAM-labeled forward primer and phosphate-labeled reverse primer, followed by λ-exonuclease digestion. The Round 6 pool and the starting library were diluted to 200 nM in binding buffer, heated for 10 minutes at 95°C, cooled on ice for 10 minutes, and then mixed with 3×10^5^ wild-type or GLUT1-null Caco-2 cells. Cells were incubated for 60 minutes with rotation at 4°C, washed twice with cold washing buffer, and the mean fluorescence intensity of each sample was measured on the Guava over 10,000 gated events excluding dead cells. Flow cytometry data were analyzed and plotted with Flojo.

### Next generation sequencing

The dsDNA from wild-type cell samples from Rounds 2, 4, 6, 8, and 10 and dsDNA from knockout-cell samples from Rounds 6 and 10 were sent to Vanderbilt Technologies for Advanced Genomics (VANTAGE) for NGS. 100 ng of gel-extracted dsDNA was sent for sequencing on an Illumina NovaSeq6000 PE150 Sequencer with an average of ~3.9×10^7^ raw paired-end reads per sample. The NGS data was analyzed using AptaSUITE software [27] and sorted based on their overall abundance and enrichment in each sample.

### Screening top aptamers from NGS

The top eight aptamers selected from NGS analysis were ordered from IDT with 5’-biotin labels. To screen these aptamers for their binding to the wild-type and GLUT1-null Caco-2 cells, the aptamer stocks were diluted to 200 nM in binding buffer, heated at 95°C for 10 minutes, cooled on ice, and mixed with 3×10^5^ cells. After 60 minutes of incubation with rotation at 4°C, the cells were washed twice with cold washing buffer. The cells were then resuspended in 1:200 dilution of streptavidin, Alexa Fluor 488 conjugate (Thermo Fisher, S11223) and incubated for 30 minutes at 4°C with rotation. The cells were washed a final two times and measured on the Guava flow cytometer for their mean fluorescence intensity over 10,000 gated events. The top four aptamers from this flow cytometry screen were order from IDT with FAM fluorophore labels, along with an equilength FAM-labeled scrambled aptamer, to perform live-cell imaging. 1×10^5^ wild-type and GLUT1-null Caco-2 cells were seeded into 24 well plates (Corning, 3526) and allowed to adhere overnight. The following day, cells were washed twice with wash buffer and blocked for 15 minutes with binding buffer at 4°C. Individual aptamers were diluted to 200 nM in binding buffer, heat prepped as described above, and incubated with cells at 4°C for 30 minutes. After incubation, the cells were incubated with the Cytopainter cell membrane stain (Abcam, ab219941), washed three times with washing buffer, and then imaged on a Leica DMi8 fluorescence microscope.

### Generation of a homogenous GLUT1 expressing Caco-2 cell line

For affinity tests, a homogenous GLUT1+ Caco-2 cell line (termed high-expressing GLUT1 Caco-2) was generated by FACS with an Alexa Fluor 488 primary conjugated GLUT1 antibody (FAB1418G, 1:1000 dilution). Briefly, Caco-2 cells were washed twice with DPBS, dissociated, and resuspended in phenol-free DMEM medium supplemented with 10 μM Y27632 and 1X antibiotic-antimycotic. Fluorescent cells were sorted into 12 well plates using the 5-laser FACS Aria III with a 100 μm nozzle. Sorted cells were supplemented with complete growth medium 24 hours after sort and medium was changed with complete growth medium every 48 hours afterwards. Despite some controversy as to whether this antibody recognizes an extracellular epitope on GLUT1 [28], [29], we verified homogenous GLUT1 expression in the sorted cells using western blot analysis, immunofluorescent imaging, and flow cytometry.

### Flow cytometry analysis of top binding aptamer

The affinity and specificity of aptamer A5 were measured by flow cytometry. To determine the affinity of the A5 aptamer and scrambled aptamer control for high-expressing and GLUT1-null Caco-2 cells, the aptamers were serially diluted from 500 nM to 0 nM in binding buffer. Cells were washed twice and dissociated with 0.02% EDTA. 3×10^5^ cells were incubated in each aptamer dilution for 1 hour at 4°C and measured for their mean fluorescence intensity on the Guava flow cytometer. Graphpad Prism was used to determine the apparent dissociation constant (K_d_) of the aptamer binding by fitting the data to a standard one-site, specific binding model. The specificity of aptamer A5 was determined by performing qualitative flow cytometry binding experiments using HEK-293 cells, MDA-MB-231 cells, and iPSC-BMECs. In these experiments, 3×10^5^ cells were incubated with 200 nM aptamer A5 in binding buffer at 4°C for 1 hour, washed twice, and measured for their mean fluorescence intensity on the Guava flow cytometer. Mean fluorescent binding values of aptamers were compared to values obtained for a GLUT1 antibody targeting an intracellular epitope (Abcam, ab15309, 1:1000 dilution). For this comparison, cells were fixed in 4% PFA for 20 minutes, blocked with PBS containing 5% donkey serum and 0.3% Triton-X-100 for 1 hour, and incubated with the primary GLUT1 antibody overnight at 4°C. The following day, cells were incubated with Alexa 488 donkey anti-rabbit secondary for 1 hour, washed three times, and analyzed for mean fluorescence intensity in the same manner.

### Tissue imaging

De-identified human brain tissue samples were provided by the Cooperative Human Tissue Network Western Division run by Vanderbilt University Medical Center. Tissue blocks were embedded in optimal cutting temperature compound (Fisher, 23730571), sliced into 10 μm sections, mounted on charged glass slides, and stored at −80°C. For imaging, the tissue sections were warmed to room temperature and blocked with binding buffer containing 20% FBS and 1 mg/ml yeast tRNA for 60 minutes. Following blocking, the tissue sections were incubated with 200 μL of 250 nM FAM-labeled aptamers in binding buffer or 200 μL of GLUT1 antibody (Abcam, ab15309, 1:1000 dilution) in PBS with 2% FBS for 1 hour on ice in the dark. For tissue sections incubated with antibody, Alexa Fluor 488 conjugated donkey anti-rabbit secondary was incubated for 1 hour following primary incubation. Tissue sections were then washed three times with washing buffer and co-stained with DAPI in PBS for 10 minutes. Tissue sections were then mounted with anti-fade reagent (Thermo Fisher, P10144) and imaged using a Leica DMi8 fluorescence microscope. For dual imaging, tissue sections were incubated with 200 μL of 250 nM Alexa Fluor 647-labeled aptamers in binding buffer or GLUT1 antibody in PBS with 2% FBS for 1 hour on ice in the dark. Antibody-incubated tissue sections were followed by treatment with Alexa Fluor 647 conjugated donkey anti-mouse secondary (Thermo Fisher, A-31573) for 1 hour. Tissue sections were then incubated with DyLight 488 labeled lycopersicon esculentum lectin (Vector Laboratories, DL-1174 1:100 dilution) in PBS for 30 minutes, followed by DAPI for 10 minutes. Tissue sections were then mounted and imaged.

### Measurements of aptamer serum stability

To determine the serum stability of aptamer A5, 1 μg of aptamer was incubated with 50% FBS at 37°C for the following time points: 0, 0.25, 0.5, 1, 2, 4, 8, 12, 24, and 30 hours. Following the incubation, the DNA was run on a 3% agarose gel and visualized using an Odyssey Fc Imager.

### Measurements of glucose uptake into cells

2×10^4^ wild-type Caco-2 cells were seeded into each well of a 96 well plate and allowed to adhere overnight. The following day, glucose uptake was measured using the Glucose Uptake-Glo Assay (Promega, J1341). Prior to performing the experiment, cells were incubated with 50 μM of a single aptamer (A1-A8) for 30 min, 50 μM of cytochalasin B (Sigma C6762) for 5 minutes, or left untreated. Cells were then washed to remove any glucose, incubated with 50 μL of 2-Deoxy-D-glucose (2-DG), and incubated with stop and neutralization buffers according to the manufacturer’s instructions. 100 μL of glucose-6-phosphate dehydrogenase (G6PDH) were added to each well, incubated for 2 hours, and then luminescence was measured on a Tecan Inifinite M1000 Pro. Glucose uptake inhibition by aptamers A1-A8 was compared to inhibition by cytochalasin B, cells that did not receive 2-DG, and untreated cells.

## Results and Discussion

### Generation of GLUT1-null Caco-2 cells

To select aptamers against the GLUT1 transporter, we first sought to engineer a GLUT1-null cell line by using CRISPR techniques to knock out the *SLC2A1* gene, which encodes the GLUT1 glucose transporter. This GLUT1-null cell line was intended for use in the negative selection step as the isogenic pair to remove any aptamers from the selection process that were not specific to the GLUT1 transporter. Wild-type cells would then be used in the positive selection step as they express GLUT1 as the primary target. The Caco-2 epithelial cell line was chosen for this aptamer selection due to its high predicted expression of GLUT1 by the Human Protein Atlas [30]. To facilitate the gene editing process, we engineered a Caco-2 line with constitutive Cas9 expression using lentiviral transduction and clonal sorting (**Fig 2A, 2B**). Immunofluorescent imaging (**Fig 2C**) and flow cytometry analysis (**SI Fig 1**) were initially used to demonstrate that the GLUT1 transporter was expressed in the Cas9-expressing Caco-2 cells, albeit heterogeneously. To generate the *SLC2A1* knockout, synthetic gRNAs were delivered to the Caco-2 cells using lipofection, and after three rounds of gRNA treatment, single clones were picked for expansion and initial analysis by western blot (data not shown). One clone was chosen for full analysis, and GLUT1 knockout in this clone was validated with immunofluorescent imaging (**Fig 2C**) and western blot (**Fig 2D**). Additionally, the online Tracking of Indels by DEcomposition (TIDE) [25] genomic assessment tool was used to quantify the percent knockout. By computational assessment of Sanger-sequenced genomic DNA, two of the four cut sites were predicted to have indel formations of at least 94% (**Fig 2E**), which is highly indicative of a knockout. Collectively, these results demonstrate the engineered Caco-2 cell line has complete knockout of the GLUT1 protein and was therefore suitable for use in our aptamer selection.

**Figure 2.**
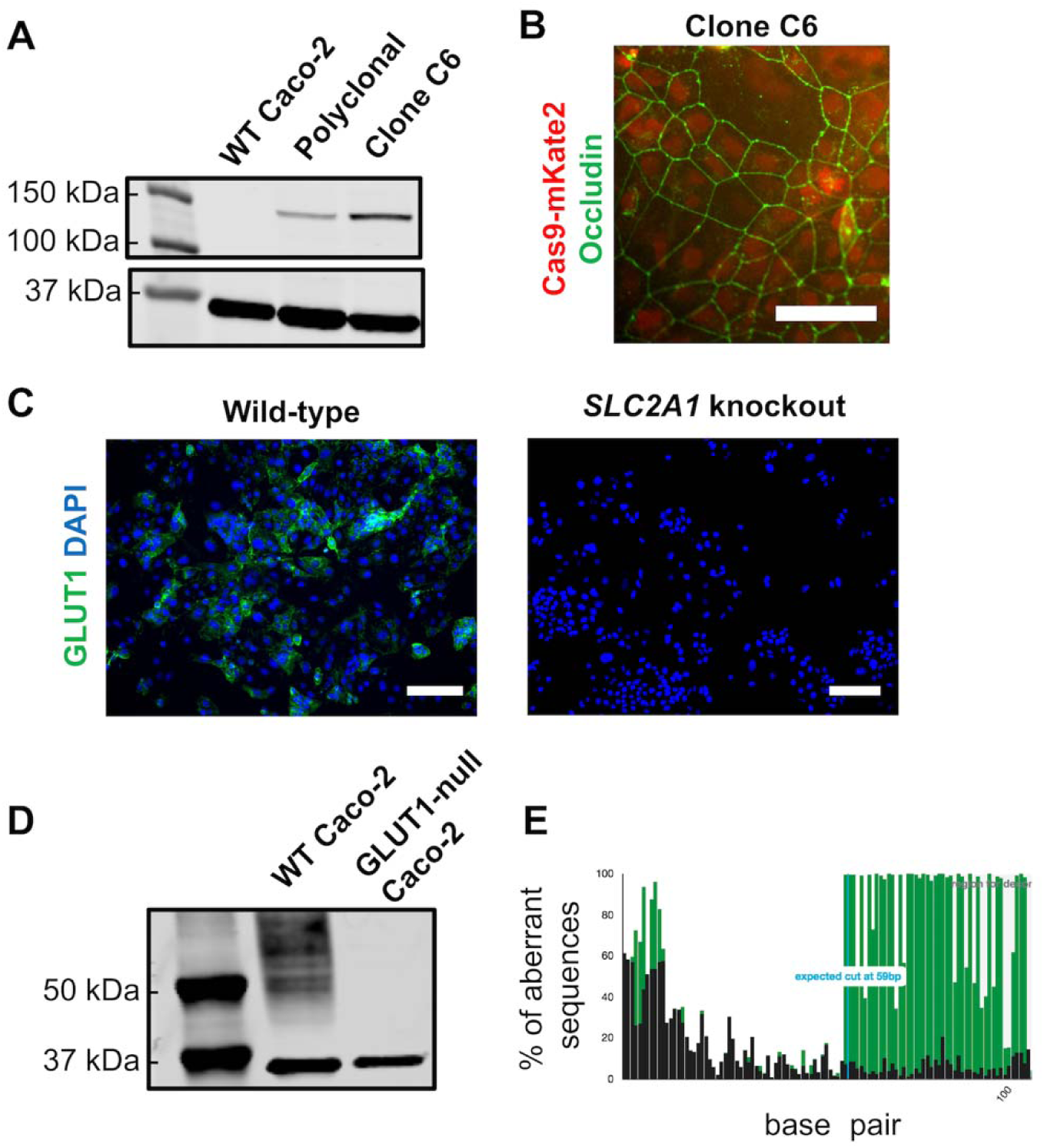
Generation of GLUT1-null Caco-2 cells. (A) Cas9-expressing Caco-2 cells were created by transduction with lentivirus and clonal isolation. Western blot analysis was used to confirm Cas9 expression in the polyclonal line and the final selected clone C6, with GAPDH as the housekeeping protein. (B) Immunofluorescent image showing Cas9 expression in clone C6, with Cas9 fused to mKate2 (red) and occludin (green) marking the cell borders (scale bar: 100 μm). (C) Immunofluorescent staining of GLUT1 in wild-type versus *SLC2A1* knockout Caco-2 cells (referred to as GLUT1-null Caco-2) (scale bar: 200 μm). (D) Western blot image of GLUT1 protein expression (expected band size of 55 kDa) in wild-type (WT) versus GLUT1-null Caco-2 cells. GAPDH was used as the housekeeping protein, with bands visible at correct band size of 37 kDa for both cell lines. (E) Example TIDE genomic assessment data for one of the four expected gRNA cut sites. The black sequence is the control sample (wild-type Caco-2 cells) and the green sequence is the test sample (*SLC2A1* knockout). The TIDE web tool predicts an indel formation of 96.1% for this particular cut site.

### Selection of DNA aptamers against GLUT1

Wild-type and GLUT1-null Caco-2 cells were used for the positive and negative selection steps of CRISPR-mediated isogenic cell-SELEX, respectively. The selection process is summarized in **Figure 1**, building upon established protocols [3], [6]. In the first round of this selection process, a random 40-mer DNA library was perfused over the wild-type Caco-2 cells to enrich the library for binding to the wild-type GLUT1+ cells. Following the first round, a negative selection step was added to the positive selection step, by perfusing the pool over the GLUT1-null cells followed by the wild-type cells. In this process, the pool is continuously circulated over both cell types for a fixed amount of time, in an effort to allow the nucleic acid pool to interact with both cell types in a single round of selection; we recently used this strategy for isolating aptamers against recombinant protein targets [26]. Following this simultaneous positive and negative selection, cells were washed and aptamers were eluted from each Caco-2 population. ssDNA was PCR amplified and the resultant dsDNA was restored to single-strand form with enzyme digestion to be introduced into the next round of selection.

After Round 6 of selection, the pool affinity was tracked with a flow cytometry binding assay. The Round 6 pool was amplified with FAM-labeled forward primer and showed preferential binding to the wild-type Caco-2 cells compared to the fluorescently-labeled starting library (**Fig 3A**). Additionally, the Round 6 pool showed negligible binding as to the GLUT1-null Caco-2 cells, similar to the starting library (**Fig 3B**). Thus, through six rounds, it appeared that the nucleic acid pool was enriching to bind the wild-type cells and not the GLUT1-null cells. Four more rounds of selection were completed to further enrich the pools. In total, ten rounds of selection were performed, with ten positive selection steps and nine negative selection steps. After completion of the selection, samples of the pools from Rounds 2, 4, 6, 8, and 10, along with samples from the negative selection from Rounds 6 and 10, were subjected to NGS analysis. Sequencing data were then analyzed with AptaSUITE bioinformatics software [27]. The amount of unique sequences was tracked in each sample, with a sharp convergence in the number of sequences after round 6 (**Fig 4A**). Sequences were ranked in each sample based on their overall abundance (the overall number the sequence appeared in the pool) and enrichment (fold-change in any particular sequence between two rounds), similarly to previous reports [31], [32]. Based on this analysis, eight prospective aptamers were selected from the NGS data (**Fig 4B**), including the three most abundant sequences from Round 10 (A1-A3), the top enriched sequence from Round 10 (A4), the top enriched sequence from Round 6 (A5), the second most abundant sequence in Round 6 (A6), the fourth most abundant sequence from Round 6 (A7), and the top enriched aptamer from Round 4 (A8). These sequences were cross-compared with the NGS data from the negative selection rounds to make sure they were not also enriched in the GLUT1-null cells (data not shown), in an effort to avoid sequences that may have been amplified by PCR bias.

**Figure 3.**
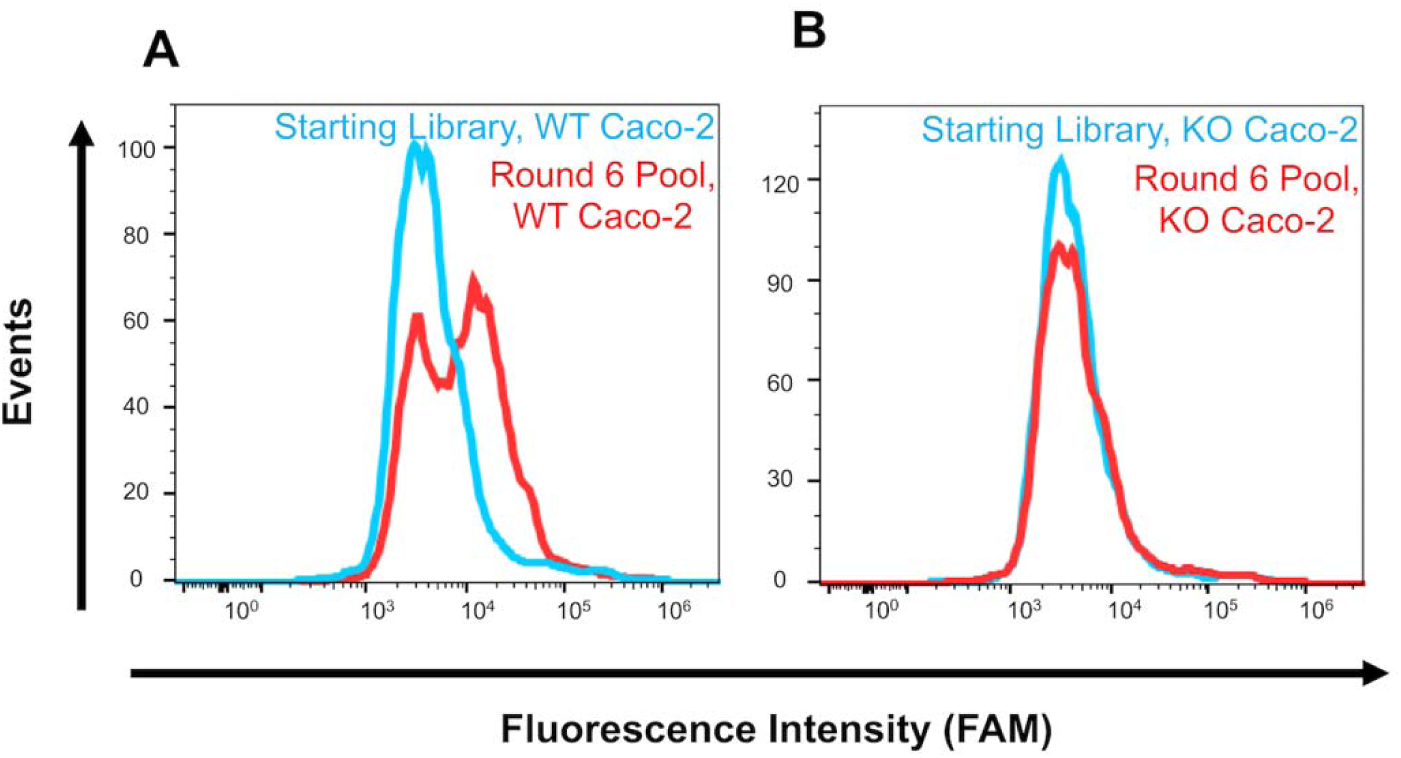
Pool affinity measurements. Affinity was measured by flow cytometry after PCR amplification of each bulk pool with FAM-labeled primers. The apparent binding of the Round 6 pool and the starting library were measured by incubating each ssDNA sample with either wild-type or GLUT1-null Caco-2 cells. (A) Histograms for binding of the Round 6 pool to the wild-type Caco-2 cells compared to the starting library. (B) Histograms for binding of the Round 6 pool to the GLUT1-null Caco-2 cells compared to the starting library. Each flow cytometry assay for the starting library and Round 6 pool was performed in duplicate against each cell type.

**Figure 4.**
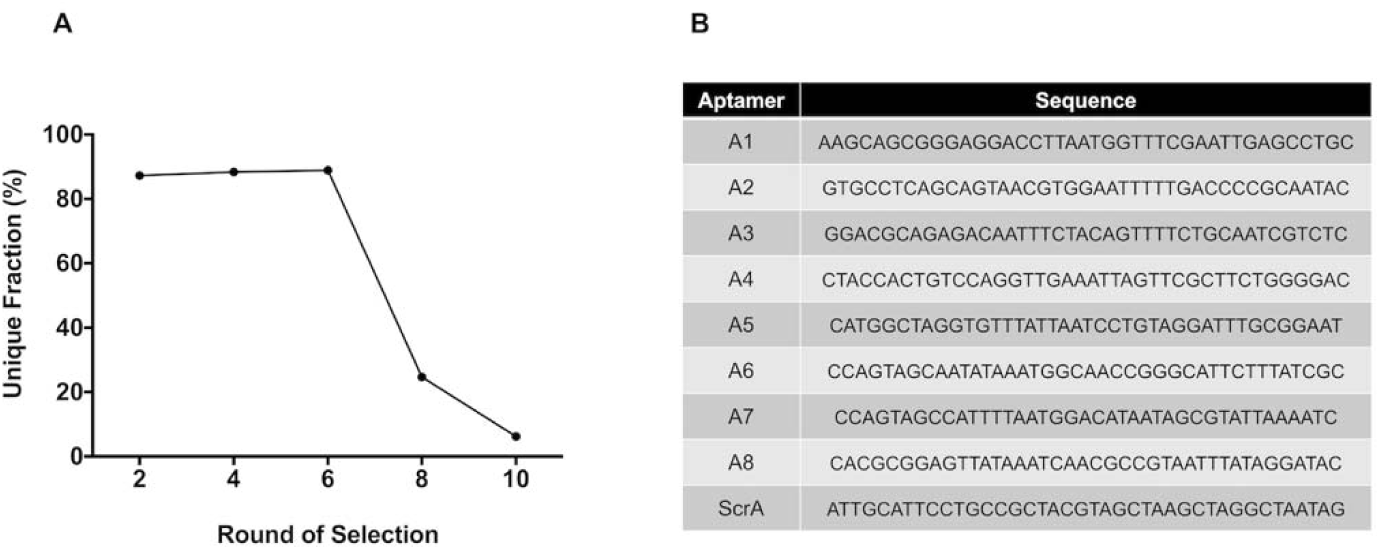
NGS analysis. NGS data was analyzed with the AptaSUITE web tool. (A) The percent of unique sequences was quantified for each of the sequenced rounds. After Round 6, the percent of unique sequences still present in the pool sharply decreases and continues to decrease through Round 10. (B) Sequences for the eight aptamers chosen for further characterization based on enrichment profiles described in the main text. The sequence for the scrambled negative control aptamer is also provided.

### Identification of GLUT1-binding aptamer candidates

The top eight sequences from the NGS data were screened for their binding to the wild-type and GLUT1-null Caco-2 cell lines using a flow cytometry assay. Aptamers A1-A8 were synthesized with biotin labels, incubated with wild-type Caco-2 cells, and mixed with streptavidin Alexa Fluor 488 conjugate, where mean fluorescence intensity was compared to a scrambled aptamer and unlabeled cells. All of the aptamers showed binding to the wild-type Caco-2 cells, compared to the controls, with several of the aptamers showing prospectively higher binding (**Fig 5A**). The mean fluorescence intensity of the aptamers was also measured for binding to the GLUT1-null Caco-2 cells, with none of the aptamers showing a strong difference in binding compared to the controls (**Fig 5A**). Because this bulk fluorescence analysis was potentially complicated by heterogeneous GLUT1 expression in the wild-type Caco-2 cells, four of the top binding aptamers were characterized further with immunofluorescent imaging. FAM-labeled aptamers A1, A2, A5, and A7 were incubated with both cell lines and live-cell images were taken and compared to the scrambled aptamer. All of the aptamers demonstrated binding to the wild-type Caco-2 cells, compared to the scrambled aptamer (**Fig 5C**). Aptamers A1, A2, and A7 showed variable adhesion to the GLUT1-null cells, likely indicating some off-target binding. However, aptamer A5 showed no apparent binding to the GLUT1-null cell line. Aptamer A5 was therefore chosen for further characterization due to its strong apparent specificity for the GLUT1 transporter. For reference, the secondary structure of aptamer A5 was predicted with NUPAC software [33] (**SI Fig 2**) and it remains stable in 50% serum for up to 12 hours (**SI Fig 3**), indicating its potential suitability for long-term imaging applications.

**Figure 5.**
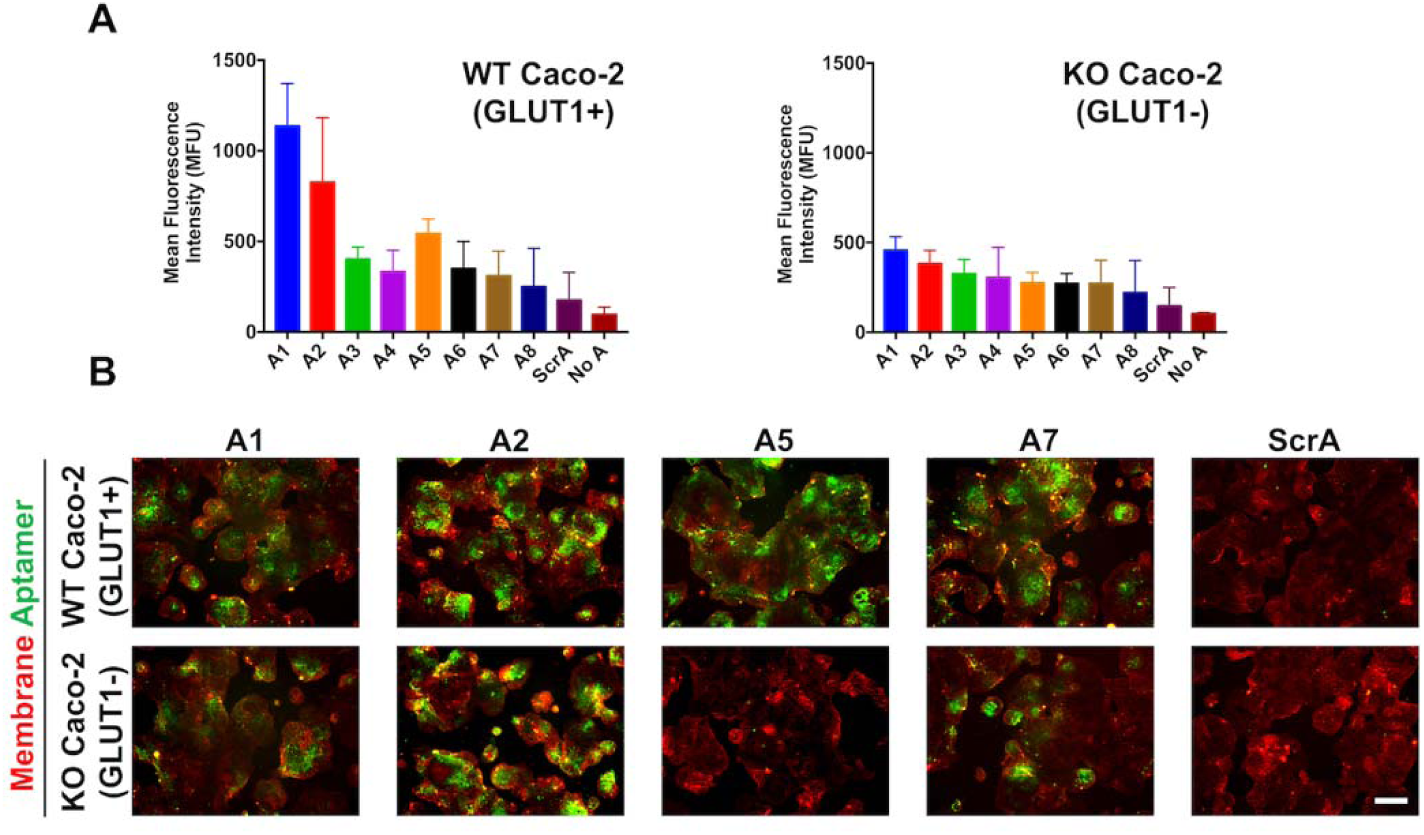
Preliminary screening of aptamer affinity and specificity. (A) Flow cytometry measurements of aptamer binding to wild-type and GLUT1-null Caco-2 cells compared to a scrambled control aptamer and unlabeled cells. Aptamers were biotin-labeled and after incubation with cells, Alexa Fluor 488-conjugated streptavidin was used for detection. Data are presented as mean ± SD from biological triplicates. (B) FAM-labeled aptamers were characterized by incubation with wild-type and GLUT1-null Caco-2 cells, followed by immunofluorescent imaging, where red indicates the cell membrane stain and green indicates the bound aptamer (scale bar: 200 μm). Images are representative of two biological replicates.

### Characterization of aptamer A5 affinity and specificity

First, to alleviate the issues encountered in **Figure 5** with a heterogeneously expressing GLUT1 population (since the presence of GLUT1-cells could skew affinity determinations), we used fluorescence-activated cell sorting (FACS) with a GLUT1-specific antibody to generate a culture of uniformly GLUT1+ Caco-2 cells (termed “high-expressing GLUT1” Caco-2 cells). This high-expressing GLUT1 Caco-2 cell line was validated by western blot, flow cytometry analysis, and immunocytochemistry (**SI Fig 4**). Flow cytometry was then used to determine the affinity of aptamer A5 to these high-expressing GLUT1 Caco-2 cells, which was measured at 160±49 nM (**Fig 6**). This result was compared to several negative controls, including a scramble aptamer incubated with the high-expressing GLUT1 Caco-2 cells and aptamer A5 incubated with the GLUT1-null Caco-2 cells (**Fig 6**). Aptamer A5 shows good selectivity as demonstrated by lower binding to the GLUT1-null Caco-2 cells (860±53 nM). The scrambled aptamer exhibits negligible binding to the high-expressing GLUT1 Caco-2 cells (e.g. binding is too low to calculate an affinity), demonstrating that these measurements are not being predominantly influenced by nonspecific interactions.

**Figure 6.**
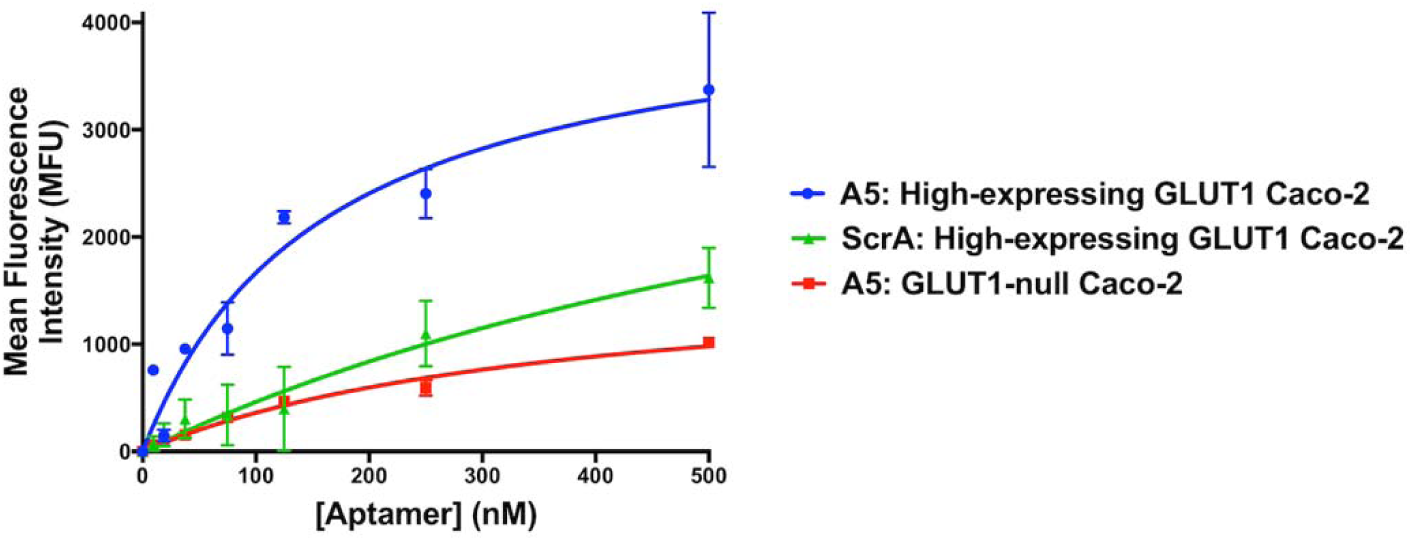
Aptamer A5 affinity analysis. Flow cytometry measurements of serially diluted (0 nM-500 nM) FAM-labeled aptamer A5 binding to high-expressing GLUT1 or GLUT1-null Caco-2 cells. The FAM-labeled scrambled aptamer was used as a negative control with the high-expressing GLUT1 Caco-2 cells. Measurements were performed in duplicate with error bars representing mean ± SD.

Beyond the quantitative comparison of binding to the GLUT1+ and GLUT1-cells, the specificity of aptamer A5 to the GLUT1 transporter was next determined by screening the aptamer against several cell lines that exhibit varying degrees of GLUT1 protein expression. The Human Protein Atlas predicts low to moderate expression of GLUT1 in HEK-293 cells and MDA-MB-231 breast cancer cells. We also routinely differentiate human induced pluripotent stem cells to brain microvascular endothelial cells (iPSC-BMECs) [23], [24], which express high levels of GLUT1. Western blot analysis was used to confirm these trends (low endogenous expression of GLUT1 in HEK-293 cells, high expression in iPSC-BMECs, and moderate expression in MDA-MB-231 cells), which were compared directly to wild-type and GLUT1-null Caco-2 cells (**Fig 7A**). Flow cytometry was then used to determine the binding of aptamer A5 to each of these cell types (**Fig 7B**). Binding patterns correlated to protein expression levels for each of the three cells types, with low binding to the HEK-293 cells, strong binding to the iPSC-BMECs, and moderate binding to the MDA-MB-231 cells. These affinity and specificity data across multiple cell types further verified the binding of aptamer A5 to the GLUT1 transporter. Additionally, the other selected aptamers GLUT1 showed similar binding patterns to aptamer A5, suggesting that while these aptamers may target other proteins, they retain some specificity for the GLUT1 transporter (**SI Fig 5**).

**Figure 7.**
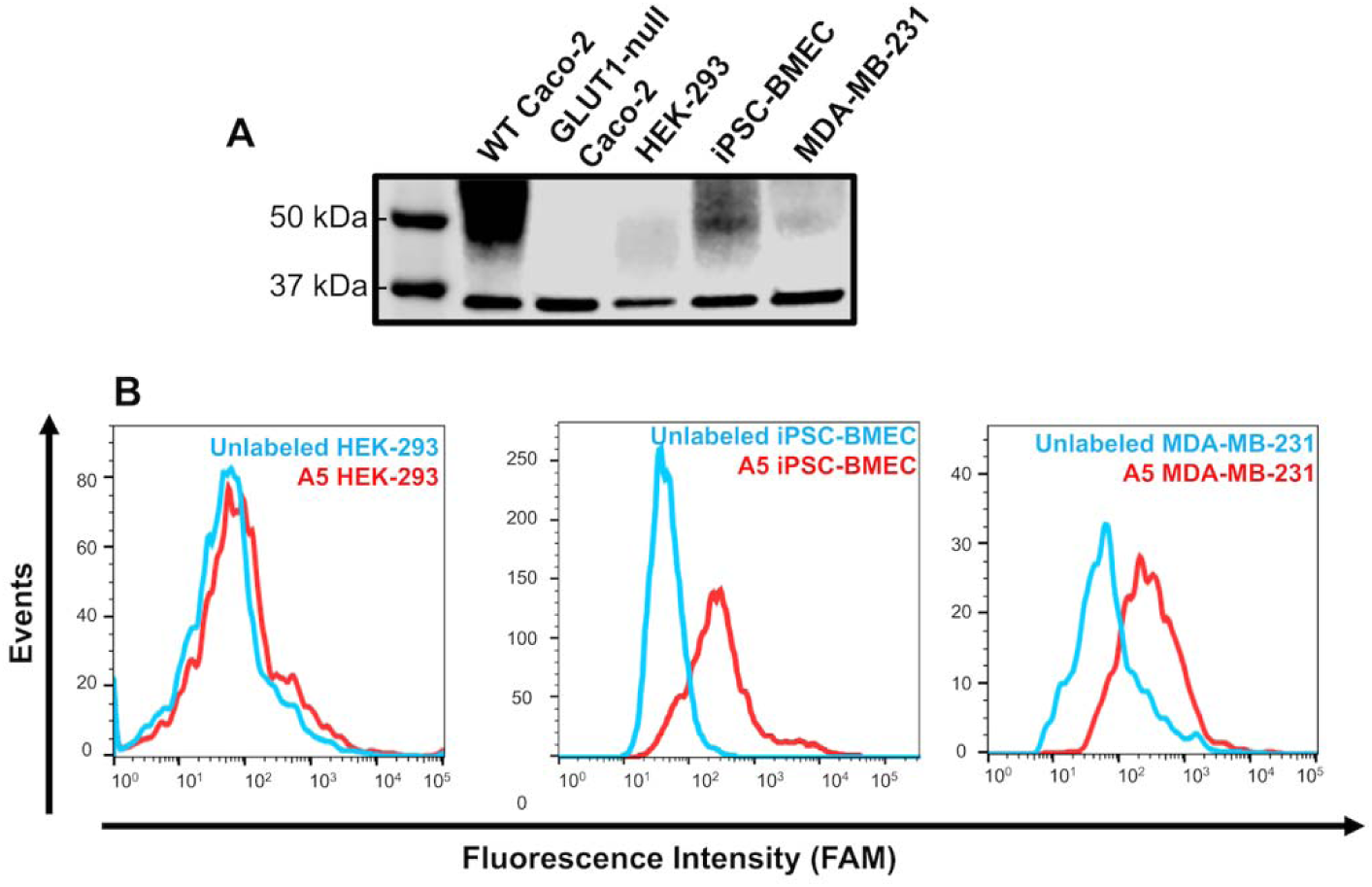
Specificity of aptamer A5 to multiple cell types. (A) Western blot showing GLUT1 expression in various cell types. GAPDH was used as the loading control. (B) Representative flow cytometry histograms for binding of FAM-labeled aptamer A5 to HEK-293 cells, iPSC-BMECs, and MDA-MB-231 cells. Binding is compared to unlabeled cells. Flow cytometry binding specificity experiments were performed in triplicate to verify trends.

### Imaging brain cortex tissue with aptamer A5

We next validated aptamer A5 using primary tissue. GLUT1 is highly expressed in brain vasculature but not in other resident neural cell types [34]. The same GLUT1 antibody used for flow cytometry analysis of cell lines showed binding to the vasculature in frozen human brain cortical tissue (**Fig 8A**), with minimal binding to other cell types. FAM-labeled aptamer A5 showed a similar binding pattern as the antibody, with minimal background and binding to other cell types (**Fig 8A**). In comparison, a FAM-labeled scrambled aptamer did not show strong binding to any cell types in the brain tissue. To further confirm the binding patterns of the aptamer to brain vasculature, the GLUT1 antibody was used in combination with a lectin dye to highlight brain vasculature (**Fig 8B**). The lectin and antibody showed similar fluorescent overlap, indicating similar binding areas. When the lectin was used in combination with an Alexa Fluor 647-conjugated aptamer A5, similar binding patterns were also observed (**Fig 8C**). These results further suggest that aptamer A5 can selectively bind the GLUT1 transporter in its native conformation.

**Figure 8.**
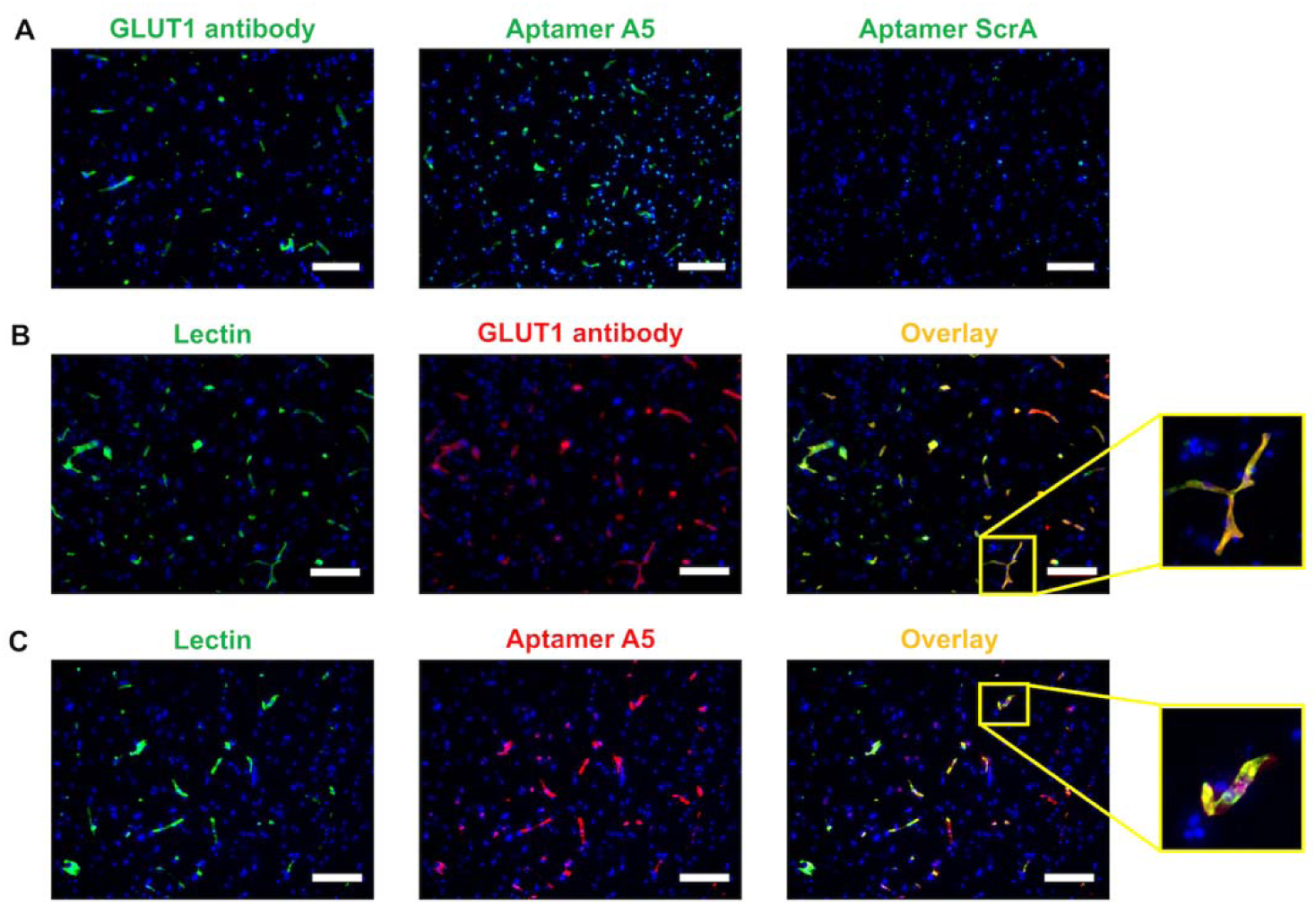
Specific binding of aptamer A5 to GLUT1 in human tissue. (A) Human brain cortical tissue was labeled with a GLUT1 Alexa Fluor 488-conjugated antibody, FAM-labeled aptamer A5, or FAM-labeled scrambled aptamer. (B) Human tissue was co-labeled with lectin and the GLUT1 antibody. (C) Human tissue was co-labeled with lectin and the Alexa Fluor 647-labeled aptamer A5. All immunolabeling was performed with two separate tissue samples, with three images taken from each tissue section to validate expression patterns. The representative images are provided with DAPI co-labeling (blue) (scale bar: 200 μm).

### Inhibiting glucose uptake with selected aptamers

Data from **Figure 5** and **SI Figure 5** indicate that other selected aptamers bind GLUT1 but lack specificity (i.e. they exhibit binding to the GLUT1-null Caco-2 cells). The Human Protein Atlas suggests that Caco-2 cells express GLUT2 and GLUT3 in addition to GLUT1, which led us to question if these nonspecific aptamers were recognizing other glucose transporters (and were thereby specific to the GLUT family, just not GLUT1 individually). As such, to further characterize these aptamers, a glucose uptake assay was performed to see if any of the eight aptamers identified with NGS could inhibit glucose entry into Caco-2 cells. Indeed, three of the aptamers (A1, A2, and A7), which also demonstrated prospective GLUT1 binding (**SI Fig 5**), yielded a significant decrease in glucose uptake that was indistinguishable from cytochalasin B, a well-characterized inhibitor of glucose transport (**SI Fig 6**). These data suggest that these aptamers might be binding to a shared epitope between GLUT transporters, without being exclusive to GLUT1. Explicit testing of GLUT2/3 knockouts would be necessary to fully validate these findings, and these experiments are being planned in the future to further qualify our cell-SELEX approach.

## Conclusion

To improve the generation of affinity reagents that bind to native cell membrane proteins with high specificity, a novel SELEX strategy was developed that utilizes isogenic cell lines. In an effort to identify aptamers specific to the GLUT1 transporter, CRISPR techniques were used to knock out the *SLC2A1* gene in Caco-2 epithelial cells to create a GLUT1-null cell line. Wild-type Caco-2 cells expressing GLUT1 were used in the positive selection step and the GLUT1-null Caco-2 cells were used in the negative selection step. Ten rounds of selection were performed and candidate aptamer sequences were identified with NGS. Most of the aptamers bound strongly to the surface of Caco-2 cells, and one of the aptamers (A5) was highly specific to the GLUT1 transporter, as demonstrated by low nanomolar affinity for wild-type but not GLUT1-null Caco-2 cells, representative binding to other cell types with varying degrees of GLUT1 expression, and selective binding to vasculature in human brain cortical tissue.

A possible limitation of this approach is that the presence of off-target CRISPR edits was not examined, so the explicit method we used to create the knockout cells could also have affected the expression of other proteins. Furthermore, although efforts were made to control parameters such as density, which can influence receptor expression levels [35], [36], fluctuations may have occurred between selection rounds to influence the final outcomes. In addition, despite the clonal selection of the original Cas9-expressing Caco-2 cells, mutations may have arisen throughout the SELEX process as the cells were continuously propagated, which could reimpose some heterogeneity outside of the GLUT1 knockout. Last, while this cell-SELEX approach can be definitively used to obtain highly specific aptamers, more in-depth cell engineering to remove homologous family members may be necessary to improve the hit rate of selected aptamers. Other users should be mindful of these variables when utilizing this cell-SELEX approach.

Overall, our data indicate that highly specific aptamers can be efficiently isolated against native membrane proteins with our SELEX strategy that uses CRISPR-engineered isogenic cell lines. This approach should be broadly useful for generating affinity reagents that bind to diverse classes of membrane proteins with high specificity. We also expect that isogenic whole cell selection methods could be extended to other classes of amino acid-based affinity reagents with similar success. Future efforts will focus on strategies for improving the affinity of selected aptamers through random or site-directed mutagenesis.

## Abbreviations

SELEX: Systematic Evolution of Ligands through Exponential Enrichment
CRISPR: Clustered Regularly Interspaced Short Palindromic Repeats
GLUT: Glucose Transporter
TIDE: Tracking of Indels by Decomposition
NGS: Next-Generation Sequencing
iPSC: Induced Pluripotent Stem Cell
BMEC: Brain Microvascular Endothelial Cells

## Acknowledgments

We gratefully acknowledge Allison Bosworth for providing iPSC-derived BMECs, Ella Hoogenboezem for providing MDA-MB-231 cells, and Everett Allchin for culturing HEK-293 cells. This research was supported by a Ben Barres Early Career Acceleration Award from the Chan Zuckerberg Initiative (grant 2018-191850 to ESL), grant A20170945 from the BrightFocus Foundation (ESL), grant IRG-58-009-56 from the American Cancer Society (ESL), and an Engineering Immunity Pilot Grant from Vanderbilt University (ESL). This work was further supported by facilities at Vanderbilt University, including the Vanderbilt University Medical Center Flow Cytometry shared resource (supported by the Vanderbilt Ingram Cancer Center NIH grant P30 CA68485 and the Vanderbilt Digestive Disease Research Center NIH grant DK058404) and the Vanderbilt Technologies for Advanced Genomics core facility (supported by the Vanderbilt Ingram Cancer Center NIH grant P30 CA68485, the Vanderbilt Vision Center NIH grant P30 EY08126, and NIH/NCRR grant G20 RR030956). EHN is supported by a Graduate Research Fellowship from the National Science Foundation (DGE-1445197). DAB was supported by the Vanderbilt University Medical Scientist Training Program (T32 GM007347). We thank the Cooperative Human Tissue Network, an NIH/NCI sponsored resource, for providing human brain tissue.

## Conflict of interest

The authors declare no conflicts of interest.

## Ethical approval

All human tissue used in this study was de-identified and did not require IRB approval.

