## Supplemental Information for "CRISPR-mediated isogenic cell-SELEX approach for generating highly specific aptamers against native membrane proteins"

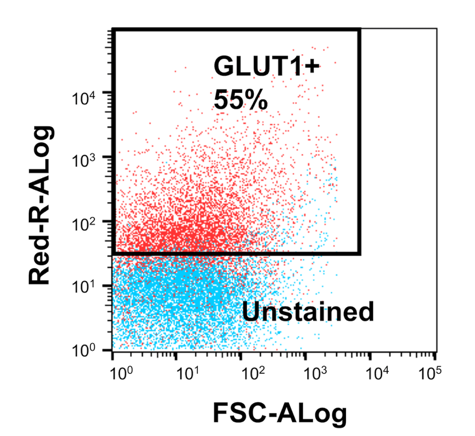

**SI Figure 1.** Flow cytometry dot plot showing relative expression of GLUT1 in wild-type Caco-2 cells using a GLUT1 antibody conjugated with Alexa Fluor 647. Caco-2 cells heterogeneously express GLUT1 (~55% of the total population) under the conditions used for SELEX.

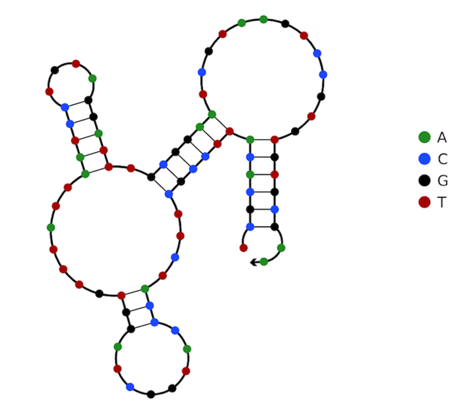

**SI Figure 2. Aptamer A5 structure.** NUPACK nucleic acid software was used to predict the structure of aptamer A5 [1].

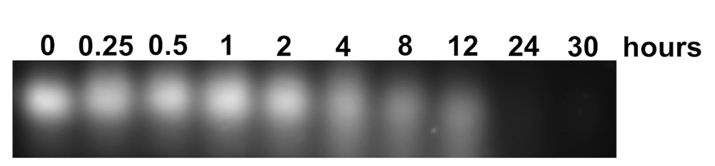

**SI Figure 3. Serum stability of aptamer A5.** The serum stability of aptamer A5 was tested by incubation with 50% serum for various time points from 0 to 30 hours. DNA samples were visualized in a 3% agarose gel.

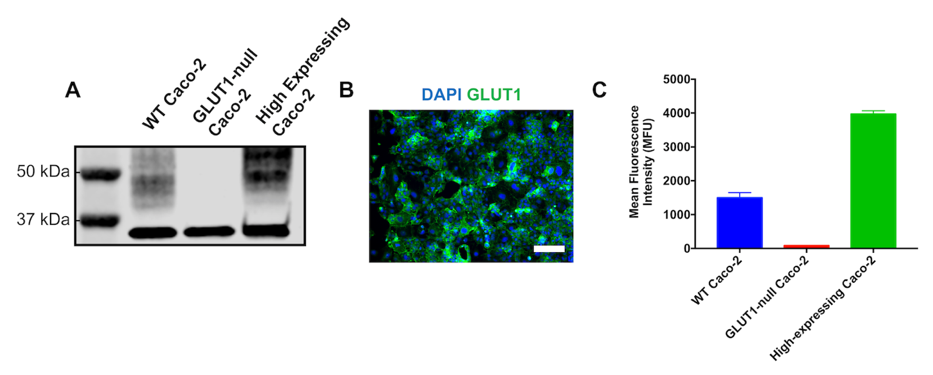

**SI Figure 4. Validation of FACS-sorted high-expressing Caco-2 cells used for affinity measurements.** Caco-2 cells were FACS-enriched with an Alexa Fluor 488-conjugated GLUT1 antibody and assessed for GLUT1 expression. (A) Western blot for GLUT1 expression. (B) Immunofluorescent imaging demonstrates relatively homogenous expression of GLUT1 (scale bar: 200 μm). (C) Flow cytometry measurements of GLUT1 antibody binding to wild-type Caco-2 cells, GLUT1-null Caco-2 cells, and high-expressing GLUT1 Caco-2 cells. Flow cytometry binding experiments performed in duplicate with error bars representing mean ± SD.

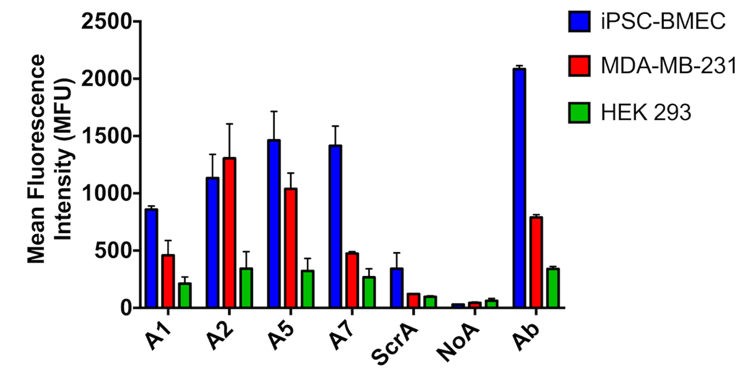

**SI Figure 5. Aptamer specificity against multiple cell types.** Aptamers A1, A2, A5, and A7 were screened against three cell types expressing different levels of GLUT1 (iPSC-BMEC, MDA-MB-231, and HEK-293). Mean aptamer fluorescence intensity was compared to a scrambled aptamer, unstained cells (noA), and a GLUT1 antibody (Ab). Flow cytometry binding experiments were performed in triplicate with error bars representing mean ± SD.

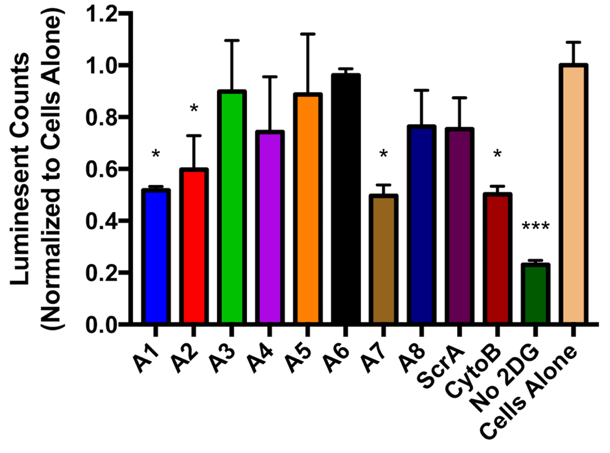

**SI Figure 6. Glucose uptake assay.** Glucose uptake was measured via 2-deoxy-D-glucose (2-DG) uptake using the Uptake-Glo assay. Glucose uptake was compared between Caco-2 cells incubated with aptamers A1-A8, the scrambled aptamer, cytochalasin B, cells that did not receive any prospective inhibitor (cells alone), and cells that did not receive any 2-DG. Luminescent counts were normalized to cells alone. The assay was performed once with duplicate wells for each condition with error bars representing mean ± SD. Significance was determined with an ordinary one-way ANOVA in Graphpad (*, p<0.06; ***, p<0.001; all other aptamers, p>0.35).
